# From ecology to evolution: plasmid- and colicin-mediated persistence of antibiotic resistant *Escherichia coli* in gulls

**DOI:** 10.1101/2025.09.30.679527

**Authors:** Michaela Ruzickova, Jana Palkovicova, Kristina Nesporova, Marketa Rysava, Rene Pariza, Simon Krejci, Ivan Literak, Monika Dolejska

## Abstract

Antimicrobial resistance in wildlife is an emerging concern within the One Health concept. Gulls, due to their synanthropic behaviour and long-distance migration, are recognised as vectors and secondary reservoirs of resistant bacteria. These birds can facilitate the environmental spread of resistant strains across ecosystem boundaries. Understanding their role in shaping microbial communities is essential for assessing the broader ecological impact. This study investigates the persistence and competitive dynamics of cephalosporin-resistant *Escherichia coli* in Caspian gulls captured at their breeding colony at a water reservoir and subsequently monitored in captivity for three months, representing the longest *in vivo* experiment of its kind conducted on wild birds.

We observed sustained colonization and long-term shedding of resistant *E. coli* throughout the entire study, marking the longest documented carriage of resistant bacteria in wild birds to date. Notably, rapid dissemination of various *E. coli* sequence types (STs) with CTX-M-1 was observed, with ST11138 rapidly outcompeting other strains, including the initially dominant ST11893. Genomic analyses revealed that ST11138 harboured F24:A-:B1 and IncI1/ST3/CTX-M-1 plasmids encoding colicins and corresponding immunity genes, likely conferring a competitive advantage.

Our findings underscore the role of bacteriocin-mediated interactions in shaping microbial communities and highlight the importance of plasmid-encoded traits in the persistence of resistant strains in wildlife. Importantly, our findings underscore the ecological novelty of longitudinal *in vivo* tracking of AMR persistence in natural hosts and highlight the need to consider ecological and microbiome-level interactions when assessing the environmental dimension of AMR under the One Health concept.

**Importance:** Antimicrobial resistance in wildlife is an emerging concern within the One Health framework, with gulls recognized as important vectors and secondary reservoirs of resistant bacteria. Due to their synanthropic behaviour and long-distance migration, these birds can facilitate the spread of resistant strains across ecosystems. However, the role of wildlife in resistance dynamics remains underexplored, especially in long-term, natural settings. Our study is unique in its scope and duration, representing the longest *in vivo* experiment of its kind conducted on wild birds. By capturing these processes in live hosts under naturalistic conditions and across an extended period, our study provides rare and ecologically grounded insights into how antimicrobial resistance is maintained outside clinical or laboratory settings. Our findings show sustained colonization and long-term shedding of resistant *E. coli*, with strain ST11138 outcompeting others. Genomic analyses reveal plasmid-encoded traits, highlighting the novel ecological and evolutionary mechanisms underlying resistance maintenance in wildlife.

## Introduction

Antimicrobial resistance (AMR) is a global concern with profound implications for human and veterinary medicine, agriculture, and environmental health^1^. While the role of clinical and agricultural settings in driving AMR is well established, the environmental dissemination of resistant bacteria is increasingly recognised as a critical component. In particular, the spread of AMR bacteria into natural ecosystems has been closely associated with synanthropic species, which thrive in human-altered environments and may acquire resistant bacterial strains through contact with anthropogenically impacted habitats^2,3^. Migratory birds, such as gulls (*Laridae*), are among the most effective biological vectors of AMR due to their long-distance movements and opportunistic foraging behaviour. These birds frequently exploit sites such as landfills, sewage outflows, and agricultural landscapes, where they encounter high densities of resistant microorganisms^3–5^. Their long-distance migratory behaviour enables the secondary introduction of these resistant bacteria into remote or previously unimpacted ecosystems^6,7^. Moreover, the prevalence of resistant bacteria in the vicinity of gull breeding sites has been shown to correlate with human population density and proximity to urban waste sources, suggesting that the extent of AMR within these environments mirrors broader anthropogenic pressures^8^.

The transmission of AMR is predominantly mediated by conjugative plasmids, which facilitate horizontal gene transfer across diverse bacterial species. This mechanism is especially prominent in Enterobacterales, where plasmid-mediated genes contribute to the dissemination of clinically relevant resistance traits^9,10^. Plasmids frequently harbour genes encoding the production of extended-spectrum beta-lactamases (ESBL) and AmpC cephalosporinases, conferring resistance to broad-spectrum beta-lactam antibiotics^10–12^. Notably, plasmid types carrying such resistance determinants have been identified in bacterial isolates from both wildlife and anthropogenically impacted environments, supporting the hypothesis that antibiotic resistance genes (ARGs) are actively transmitted into natural ecosystems^13–15^.

Previous studies on avian hosts have demonstrated that plasmids carrying ARGs can persist within *Enterobacteriaceae* even in the absence of direct antimicrobial selective pressure^14,16^. ESBL-producing *Escherichia coli*, in particular, exhibits the ability to survive in the avian gut for extended periods, facilitating environmental shedding and inter-individual transmission among birds as a potential strategy for maintaining colonization^14^. Although the exact duration is not established and varies strongly among bacterial species and genotypes, available studies indicate that resistant bacteria can persist in the gut of wild bird for several months^3,8^. Variation in the persistence of specific *E. coli* sequence types (STs) suggests a disparity in isolate fitness, potentially influenced by the presence of plasmids and their associated fitness costs^16,17^. Plasmids confer evolutionary advantages to STs, such as the ability to produce various bacteriocins, enabling them to outcompete other strains^18^.

In this study, we investigated cephalosporin-resistant *E. coli* isolated from the gastrointestinal tract of wild Caspian gull (*Larus cachinnans*) nestlings from a breeding colony at the Nové Mlýny water reservoir in the Czech Republic. Our primary objective was to examine the persistence and colonization dynamics of resistant strains following the removal of potential environmental sources of resistance. A longitudinal experiment including five gulls housed in controlled captivity, where individuals were sampled periodically over time, was performed. This design enabled us to monitor temporal shifts in *E. coli* STs occurrence and assess their stability. We investigated evolutionary mechanisms potentially underlying the persistence of specific STs through short- and long-read sequencing and phylogenetic analyses. When combined with other employed methods, such as conjugation frequencies, strain fitness, and phenotype analyses, these approaches provided a comprehensive insight into the factors driving strain dynamics. Particular emphasis was placed on the role of colicin, which was encoded by multiple plasmids, including the rapidly disseminating IncI1 plasmid. This plasmid, together with the colicin production, appeared to contribute significantly to the competitive success of certain strains. The study provides novel insights into the dynamics of avian gut colonization by resistant bacteria, examining persistence over a timescale that has not been previously explored. By extending the observation period beyond earlier research focused on wildlife, it sheds light on how long resistant strains may persist within wild bird populations and highlights the potential role of these hosts as long-term reservoirs.

## Material and Methods

### Sample collection

Cloacal swabs were collected in 2018 from Caspian gulls from a breeding colony inhabiting a small island in the Věstonice region of the Nové Mlýny reservoir (48.895261° N, 16.600583° E), as part of a broader telemetry study conducted during routine bird ringing^19^. This work follows a previous investigation that focused on plasmid-mediated resistance in *Escherichia* in gulls^5^. The sampling site was selected based on several factors: i) the central section of the reservoir is designated as a nature reserve and supports a diverse avian population; ii) the area is directly connected to the remainder of the reservoir, which is primarily used for recreational purposes and therefore subjected to anthropogenic influence; and iii) two major rivers flow into the central basin, one of which originates in Brno, the second largest city in the Czech Republic, where municipal wastewater, including hospital effluent, is released into the river system.

Following the initial sampling at the gull colony as part of our previous study^5^, five individuals were housed in a wildlife avian facility until they were ready to be released into the wild, equipped with the telemetry loggers^19^. Initially, each bird was kept in a separate cage; after a week, they were transferred to a shared aviary and fed a standardised diet. The selected gulls were randomly chosen non-flying juveniles, approximately one month old at the time of capture. Cloacal swabs were collected once at the gulls’ breeding site and subsequently at two-week intervals in the avian facility to monitor longitudinal changes in the resistant bacterial populations. Swabs were placed in AMIES transport medium and cultivated in buffered peptone water. Each bird was sampled on seven occasions, resulting in a total of 35 samples over the course of the study.

### Selective cultivation of resistant *E. coli*

Selective cultivation of peptone water cultures was performed on MacConkey agar (Oxoid Ltd., UK) supplemented with cefotaxime (2 mg/L). From each plate, five to six colonies with morphology consistent with *E. coli* were selected for further analysis. This approach yielded 116 isolates, of which 94 were confirmed as *E. coli* by MALDI-TOF mass spectrometry and subsequently used for further typing.

### Beta-lactamase detection

All *E. coli* isolates (n = 94) were tested by PCR for the presence of selected beta-lactamase genes, including *bla*_TEM_, *bla*_CMY-2_, and *bla*_CTX-M_. Target genes were chosen based on a previous study^5^, in which they were identified as the most prevalent among resistant *E. coli* strains from the same colony. Primer sequences are provided in Supplementary table S1.

### Pulsed-field gel electrophoresis and whole-genome sequencing (WGS)

Isolates selected based on PCR results and genomic relatedness, determined by PFGE^20,21^, were subjected to short-read WGS. PFGE macrorestriction patterns were analysed using BioNumerics v6.6 software (Applied Maths, Belgium) and are presented in Supplementary figure S1. Isolates were grouped into clusters based on 90% similarity, and representative isolates (n = 62) from each cluster were selected for further genomic analysis. Genomic DNA for short-read sequencing was extracted using NucleoSpin® Tissue kit (Macherey-Nagel, Germany). DNA libraries were prepared with the Nextera® XT Library Preparation kit (Illumina, USA) and sequenced on the Illumina NovaSeq 6000 platform run in a paired-end mode. Both the DNA library preparation and the sequencing itself was outsourced to the UTS Next Generation Sequencing Facility, Sydney, Australia.

Representative isolates (n = 4) were selected for long-read sequencing to determine the location of beta-lactamase and colicin genes and to obtain complete plasmid sequences. Isolate selection was based on short-read sequencing results and focused on STs of greatest interest: ST11138 and ST165, which were the most common STs carrying an IncI1 plasmid; ST665, the only ST harbouring an IncI1 plasmid during the first sampling; and ST11893, which carried only an IncF plasmid and was no longer detected as the experiment progressed. Genomic DNA was extracted using GenFind V3 kit (Beckman Coulter, USA). Libraries were prepared with the Rapid Barcoding Kit SQK-RBK004 (Oxford Nanopore Technologies, GB), loaded onto a FLO-MIN106 flow cell, and sequenced on the MinION Mk1B platform (Oxford Nanopore Technologies).

### Data analysis and comparative genomics

Raw reads obtained from Illumina sequencing were trimmed using Trimmomatic v0.39^22^ to remove adaptor sequences and low-quality read regions (Q ≤ 20). *De novo* assembly was performed with SPAdes v3.13.1 using the “--careful” option. Plasmid replicons, plasmid STs, and ARGs were identified using tools from the Center for Genomic Epidemiology: PlasmidFinder v2.1, pMLST v2.0, and ResFinder v4.0, respectively (https://cge.cbs.dtu.dk/services/). Colicin and associated immunity genes were assessed using ABRicate v1.0.1 (https://github.com/tseemann/abricate) with a custom database of genes generated from diverse available databases (VFDB core dataset 2025-04-15^23^, VirulenceFinder 2022-12-02^24^, and Bakta 6.0.0^25^.

Raw long reads obtained from Nanopore sequencing were demultiplexed, quality- and adaptor trimmed as described previously^26^. The processed reads were *de novo* assembled using Unicycler v0.5.1^27^ and polished by Racon v1.5.0^28^ and medaka v2.0.0^29^ using the processed long reads and by Pilon v1.2.3 with trimmed short reads. Plasmid sequences of interest were annotated by Bakta v1.10.4^25^ followed by manual curation in Geneious Prime® 2024.0.7 (http://www.geneious.com/). Genetic context of colicin and colicin immunity genes carried by plasmids was visualised using clinker v0.0.31^30^ and custom R^31^ scripts utilizing following packages: tidyverse v2.0.0^32^, ggplot2 v3.5.1^33^, dplyr v1.1.4^34^, and reshape2 v1.4.4^35^.

Plasmids were compared by BRIG v0.95^36^ using short-read assembled genomes mapped to a long-read assembled annotated reference of IncI1 and IncF plasmids. The differences among IncF and IncI1 plasmids from all isolates were assessed using alignment by Snippy v4.6.0 (https://github.com/tseemann/snippy) with corrected short-read data and long-read assembled plasmids as a reference and the nucleotide similarity was evaluated by snp-dists v0.6.3 (https://github.com/tseemann/snp-dists). The differences among IncI1 were further analysed using Varscan v2.4.6^37^.

Phylogenetic relatedness among isolates was assessed by predicting open reading frames with Prokka v1.14.1^38^, followed by core genome alignment using Roary v3.12.0^39^. This alignment served as the basis for constructing a maximum likelihood phylogenetic tree in RAxML v8.2.11^40^ under GTR+GAMMA, model supported by 1,000 bootstraps. Nucleotide similarity between isolates was estimated with snp-dists v0.6.3 (https://github.com/tseemann/snp-dists), based on the number of SNPs in the core genome alignment. The resulting phylogenetic tree was visualised in iTOL v5.7^41^.

### Competition experiments

Overnight cultures of one representative isolate of each ST11893 and ST11138 were grown separately in Luria-Bertani (LB) broth (Sigma-Aldrich, USA) at 37°C. The cultures were then diluted to 10^-3^ and incubated at 37°C until reaching an optical density at 600 nm (OD_600_) of 0.6. Equal volumes were subsequently mixed in a 1:1 ratio, while unmixed controls were maintained in parallel. All cultures were incubated for 22 hours at 37°C. Following incubation, cultures were serially diluted to 10^-7^ and plated in drops onto LB agar supplemented with cefotaxime. After overnight incubation at 37°C, colony PCR was performed on all resulting colonies to determine the relative abundance of the original ST11893 and ST11138 strains, as well as their respective transconjugants (ST11893T and ST11138T).

Custom primers targeting chromosomal markers of both STs were designed using Geneious Prime 2023.1.2. The presence of IncF and IncI1 plasmids was evaluated using primers detecting genes *bla*_CMY-2_ and *bla*_CTX-M_, which confer beta-lactam resistance and are carried by the respective plasmids. Primer sequences are provided in Supplementary table S1.

### Fitness measurements

Isolates of ST11138 (n = 6) and ST165 (n = 4), both carrying an IncI1 plasmid, were selected for growth curve analysis^9^ to compare their fitness, as they exhibited differing persistence throughout the experiment. Selection was based on both the individual host bird and the sampling time point, to capture potential diversity and temporal changes in fitness. Two isolates of ST665 were included, as this was the only ST harbouring an IncI1 plasmid at the beginning of the experiment. Additionally, two ST11893 isolates were assessed, as they carried only an IncF plasmid and disappeared during the early stages of the experiment.

Isolates were incubated for 24 hours at 37 °C, with OD_600_ measured every ten minutes. Growth curves were statistically analysed to estimate bacterial fitness. Optical density measurements were recorded using Synergy HT and Synergy HTX plate readers (BioTek Instruments, Inc., USA).

### Frequency of plasmid transfer

The same *E. coli* isolates used in the fitness assay, carrying either IncI1 or IncF plasmids (n = 14) were selected as donor strains for conjugation experiments. The recipient strain was laboratory *E. coli* MT102^42^, resistant to sodium azide (200 mg/L) and rifampicin (25 mg/L). Plasmid transfer frequency was assessed using existing protocol for filter mating assays^43^. Conjugation was performed in both technical and biological triplicates to quantify the transfer rates.

### Colicin production testing

Colicin production was assessed in the same 14 isolates used for fitness and plasmid transfer frequency experiments. Six indicator strains were utilised to cover the most commonly produced colicin types: *E. coli* K12-Row, C6 (ϕ), B1, P400 and 5K, and *Shigella sonnei* 17. The assay followed established protocols^44,45^, with *E. coli* ATCC, A15, and DH5α included as negative controls.

### Statistical analysis

Statistical computing was carried out in RStudio^46^ using custom R^31^ scripts and the following packages: readr v2.1.5^47^, magrittr v2.0.3^48^, dplyr v2.4.0^34^, and tidyverse v2.0.0^32^. Data were tested for normality using the Shapiro-Wilk test, followed by the two-tailed Mann-Whitney U test for non-parametric datasets. Significant differences were determined using a *p*-value threshold of α = 0.05. Full statistical results, including test selection and raw data, are provided in Supplementary table S2.

## Results

### High variability and dynamics of isolates

A total of 35 samples of cloacal swabs from five birds were collected, yielding 116 cephalosporin-resistant isolates, of which 94 were identified as *E. coli* by MALDI-TOF. From these, 62 isolates were selected for WGS based on pulse-field gel electrophoresis (PFGE) clustering, as well as the bird and time of sampling, in order to cover a broader range of diversity.

Across the dataset, eight different STs were identified (Fig. 1). For isolates that were not sequenced, ST assignment was inferred based on PFGE similarity to sequenced representatives. Shifts in population dynamics as well as transmission of resistance plasmids were observed during the entire experiment. At the beginning (week 0, upon admission to the avian facility), three of the five gulls carried cefotaxime-resistant *E. coli*. The most dominant strain at this time was CMY-2-producing ST11893, accounting for 81.8% (9/11) of the colonies obtained. In addition, only two ESBL-producing ST665 isolates (18.2%; 2/11) were identified in a single bird. These ST665 isolates represented the earliest carriers of the IncI1/ST3/CTX-M-1 plasmid, which subsequently disseminated into other STs.

**Figure 1.**
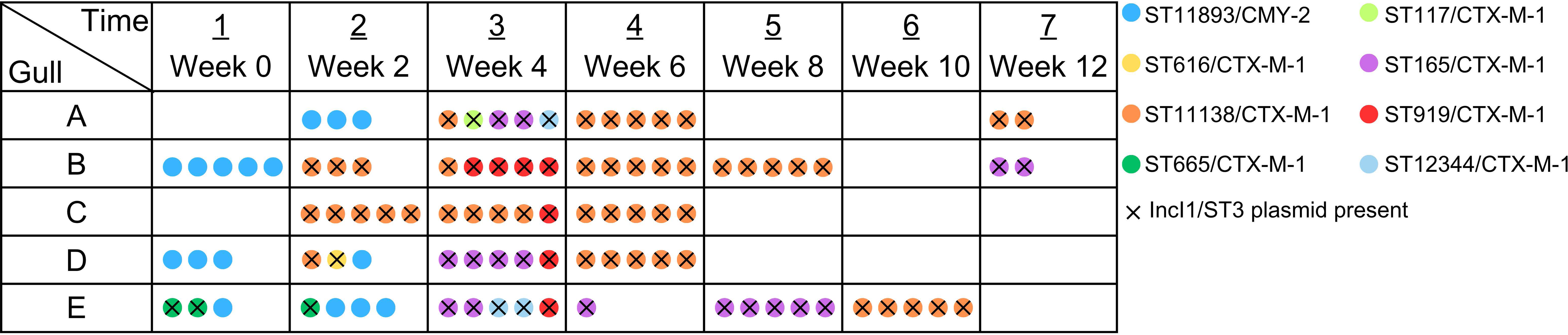
Dynamics and transmission of resistant E. coli STs in the gastrointestinal tract of gulls. Each row stands for one gull, each column for one sampling. The coloured dots represent E. coli colonies of various STs. Colonies carrying IncI1/ST3 plasmid are marked with an × symbol.

ST11893 was no longer detected after the second sampling at week 2, having been outcompeted by CTX-M-1-producing isolates carrying the IncI1/ST3 plasmid. The highest ST diversity was observed in week 4, when all birds harboured multiple STs, five in total. In the following weeks, ST11138 became dominant in all individuals, representing 50% (47/94) of the *E. coli* isolates collected. Colonisation rates declined significantly by the end of the experiment. In week 10, only one bird carried resistant *E. coli*, and this increased to two birds by week 12. Strain transmission between individual gulls was observed throughout the study period.

Analysis of phylogenetic relatedness revealed no evident branching pattern associated with individual birds or sampling time points. Instead, isolates clustered consistently according to their STs (see Supplementary figure S2). Relatively high diversity was observed even among isolates belonging to the same ST, with the highest variability found in ST11138, which displayed an average difference of 228.46 single-nucleotide polymorphisms (SNPs), and the lowest in ST11893 with an average difference of 5.04 SNPs. SNP matrices for ST11138, ST11893, ST165, and ST665 are provided in the Supplementary table S3.

### Phenotypic and molecular traits associated with strain persistence

A single ST, ST11138, repeatedly outcompeted others during the course of the experiment, suggesting a potential selective advantage. To investigate the underlying mechanisms of this dominance, four representative STs were subjected to a set of comparative assays. These included competition experiments, assessments of growth fitness, measurement of plasmid conjugation frequencies, and quantification of colicin production (Fig. 2). These analyses aimed to determine whether the observed advantage was linked to competitive ability, enhanced intrinsic fitness, or increased plasmid transfer frequency. Raw data and *p*-values for all of the analyses are provided in the Supplementary table S2.

**Figure 2.**
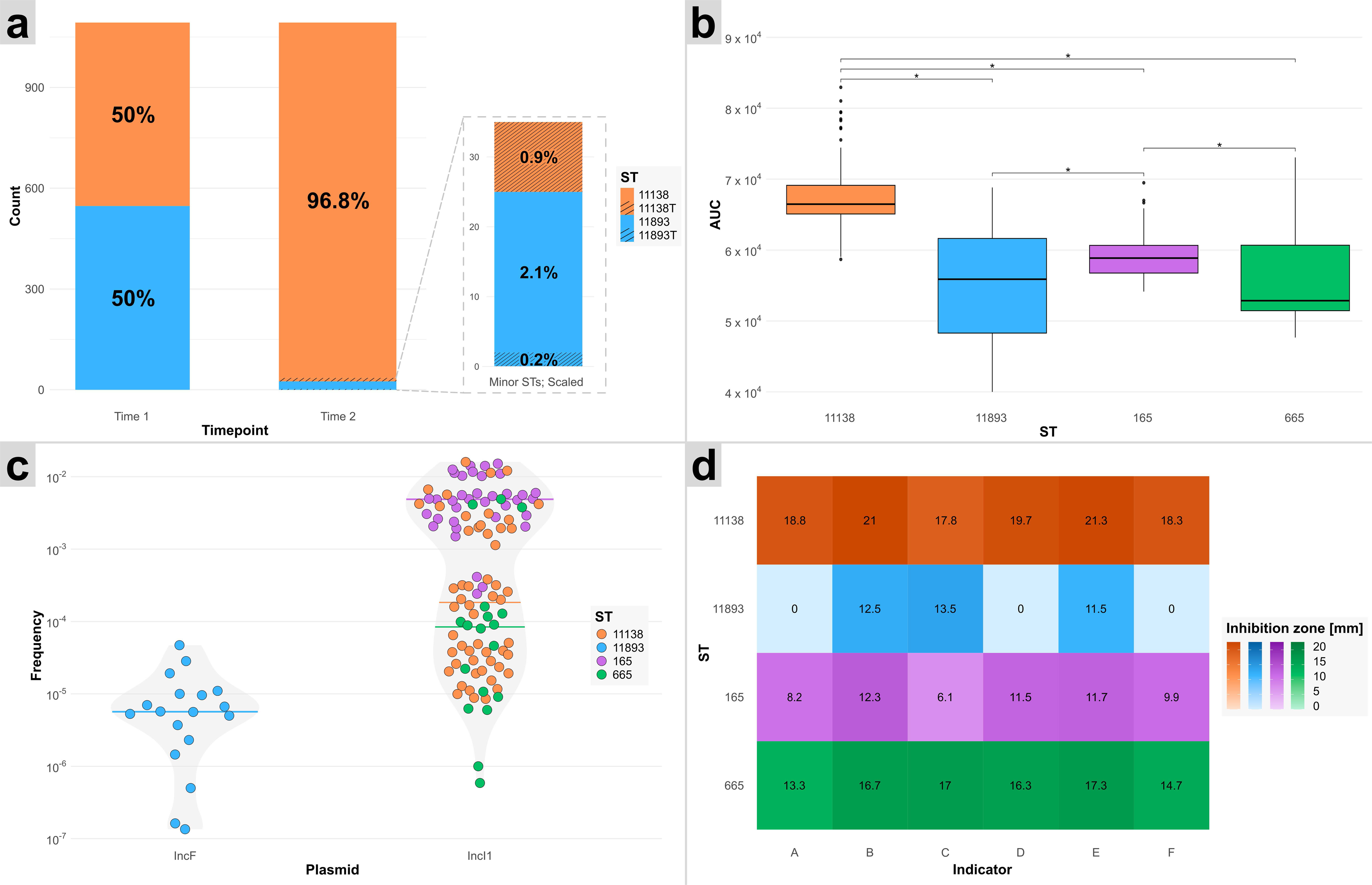
Overview of key comparative assays. (a) Results of the competition assays between ST11138 and ST11893, with minor STs enlarged to scale. The ‘T’ in ST11138T and ST11893T describes transconjugant strains. (b) Fitness of ST11138, ST11893, ST165, and ST665 based on growth curve analysis. (c) Conjugation frequencies of IncF and IncI1 plasmids, grouped by ST, with median values indicated by coloured lines. (d) Colicin production profiles, assessed by inhibition of the following indicator strains: A – E. coli 5K, B – E. coli B1, C – E. coli C6, D – E. coli K12-row, E – E. coli P400, and F – Shigella sonnei 17.

#### Competitive dominance of a single ST in competition experiments

The experiment sought to confirm the dominance of ST11138 over ST11893 under controlled laboratory conditions. Competition experiments were conducted to simulate potential interactions between *E. coli* strains within the gull host environment. Two isolates of ST11893 and six isolates of ST11138 were selected for the assay. The primary aim was to assess the dynamics within mixed populations of one ST11893 and one ST11138 isolate, including the potential transfer of the IncF plasmid from ST11893 to ST11138, and of the IncI1 plasmid from ST11138 to ST11893.

Competition assays resulted in a total number of 1,093 colonies analysed by colony PCR. Of these, 96.8% (1058/1093) were identified as ST11138 a 2.1% (23/1093) as ST11893. The rest (1.1%) displayed a conjugative transfer with 0.9% (10/1093) of ST11138 isolates obtaining the IncF plasmid (ST11138T) and 0.2% (2/1093) of ST11893 isolates obtaining the IncI1 plasmid (ST11893T) (Fig. 2a). A significant difference (*p*-value < 0.05) was observed between the final counts of most colony types, except between ST11138T and ST11893T, with ST11138 occurring at a notably higher frequency.

#### Significant differences in fitness detected among most STs

Fitness comparisons revealed significant differences between all STs, with the exception of ST11893 and ST665. Fitness was assessed using area under the curve (AUC) values derived from growth curve data. The highest fitness was observed in ST11138 (AUC 67,346.73) followed by ST165 (AUC 59,167.85), ST665 (AUC 56,602.20), and ST11893 (AUC 55,231.99), which correlated with the *in vivo* observations in live hosts (Fig. 2b).

#### Conjugation frequency of IncI1 plasmid higher than of IncF plasmid

A significant difference in conjugation rates was observed between IncI1 and IncF plasmids, with IncI1 being transferred at a 220-fold higher frequency than IncF (*p*-value = 0.03318; α = 0.05), explaining a massive IncI1 plasmid spread among all isolates (Fig. 2c). Significant differences in plasmid transfer frequencies were also detected between most STs, with the exception of comparisons involving ST665.

### Dominance of certain STs linked to variability in colicin production

Pairwise comparisons revealed statistically significant differences (*p*-value < 0.05) between multiple STs. ST11893 consistently exhibited low or no colicin production, showing significant differences from other STs across all indicator strains. In contrast, ST11138 displayed the highest levels of colicin production, differing significantly from ST165 and ST11893 across all indicators, and from ST665 specifically for indicator strains B1 and 5K.

### Varying ecological success linked to plasmid diversity and colicin genes

All sequenced isolates (n = 62) carried an IncF plasmid, with the replicon sequence type (RST) specific to the corresponding *E. coli* ST. The most common RST was F24:A-:B1 (53.2%; 33/62), found in all ST11138 and ST616. It was followed by F34:A-:B-(17.7%; 11/62) in ST11893, F-:A-:B73 (11.3%; 7/62) in ST165, F2:A-:B1 (9.7%; 6/62) in ST117 and ST919, F18:A-:B1 (4.8%; 3/62) in ST12344, and F89:A-:B-(3.2%; 2/62) in ST665. A total of 82.3% (51/62) of the isolates, all belonging to STs other than ST11893, also carried an IncI1/ST3 plasmid.

IncF plasmids of various RSTs, as well as the IncI1/ST3 plasmid, carried different combinations of five colicin production genes (*cba, cia, cib, cma, cvaC*) and their respective colicin immunity genes (c*bi, cia-imm, cib-imm, cmi, cvi*) (Fig. 3). All production genes encode eponymous colicins. The highest number of complete colicin genes was observed in ST11138, which carried *cib* on IncI1/ST3 and *cvaC* on the F24:A-:B1 plasmid, and ST165, harbouring *cib* on IncI1/ST3 and *cma* on F-:A-:B73. ST11893 carried *cma* on F34:A-:B-, and ST665 harboured *cib* on IncI1/ST3. Interrupted colicin genes were detected in ST11138 (*cia* on F24:A-:B1, disrupted by a transposase gene) and in ST11893 (*cba* on F34:A-:B-, disrupted by the resistance gene *bla*_CMY-2_). Incomplete genes were found in ST11893 (*cia* on F34:A-:B-) and ST165 (*cba* and *cvaC* on F-:A-:B73). The *cvaB* and *mchE* genes were consistently co-located with the *cvaC* on F24:A-:B1 plasmids as components of a colicin-producing operon. All STs carrying colicin production genes, whether complete, interrupted, or incomplete, also carried the corresponding immunity gene.

**Figure 3.**
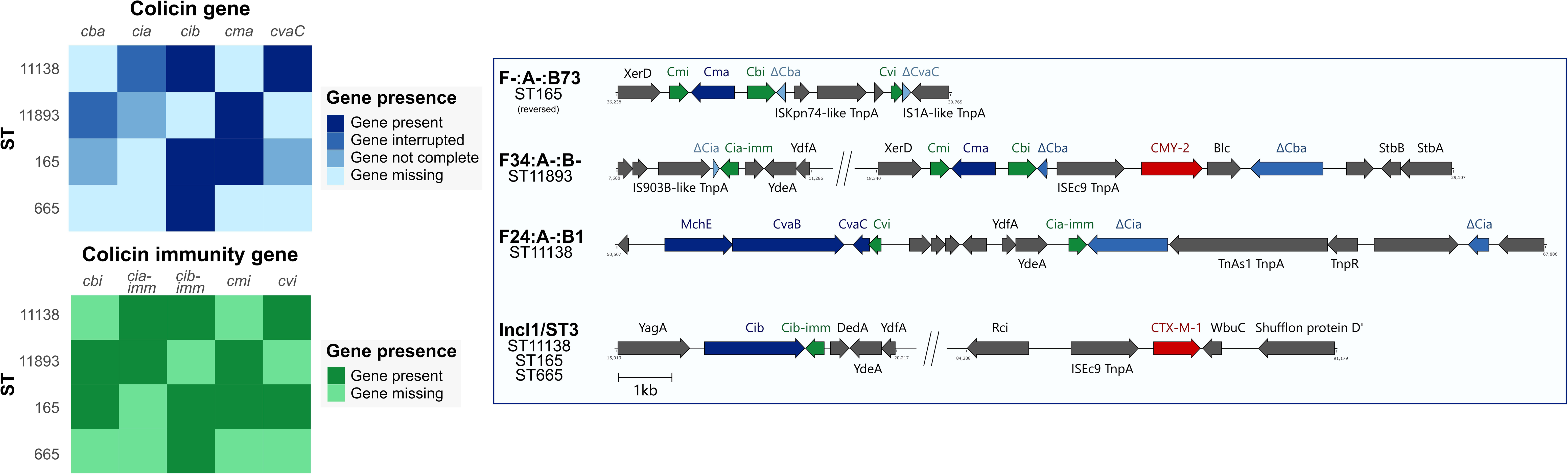
Distribution of colicin and colicin immunity genes across STs and associated plasmids. Heatmaps (left) indicate the presence, disruption, incompleteness, or absence of colicin and immunity genes across different STs. Sequences (right) show distribution of these genes across plasmids and their corresponding STs. Genes encoding colicin production are highlighted in blue, immunity genes in green, and antibiotic resistance genes in red.

F24:A-:B1 plasmid carried by the ST616 isolate differed distinctly from those found in ST11138 isolates, which exhibited high sequence uniformity (Fig. 4). Notable differences were observed in the presence of several colicin genes (*cia, cia-imm*, *cvaC*) as well as ARGs (*macA*, *macB*). SNP analysis confirmed the uniformity of F24:A-:B1 plasmid among ST11138 isolates with no SNPs detected between them. In comparison, the variant of the same plasmid harboured by ST616 differed by 884 SNPs when aligned against all other sequences (Supplementary table S4). For a complete alignment of the F24:A-:B1 plasmids, see Supplementary figure S3.

**Figure 4.**
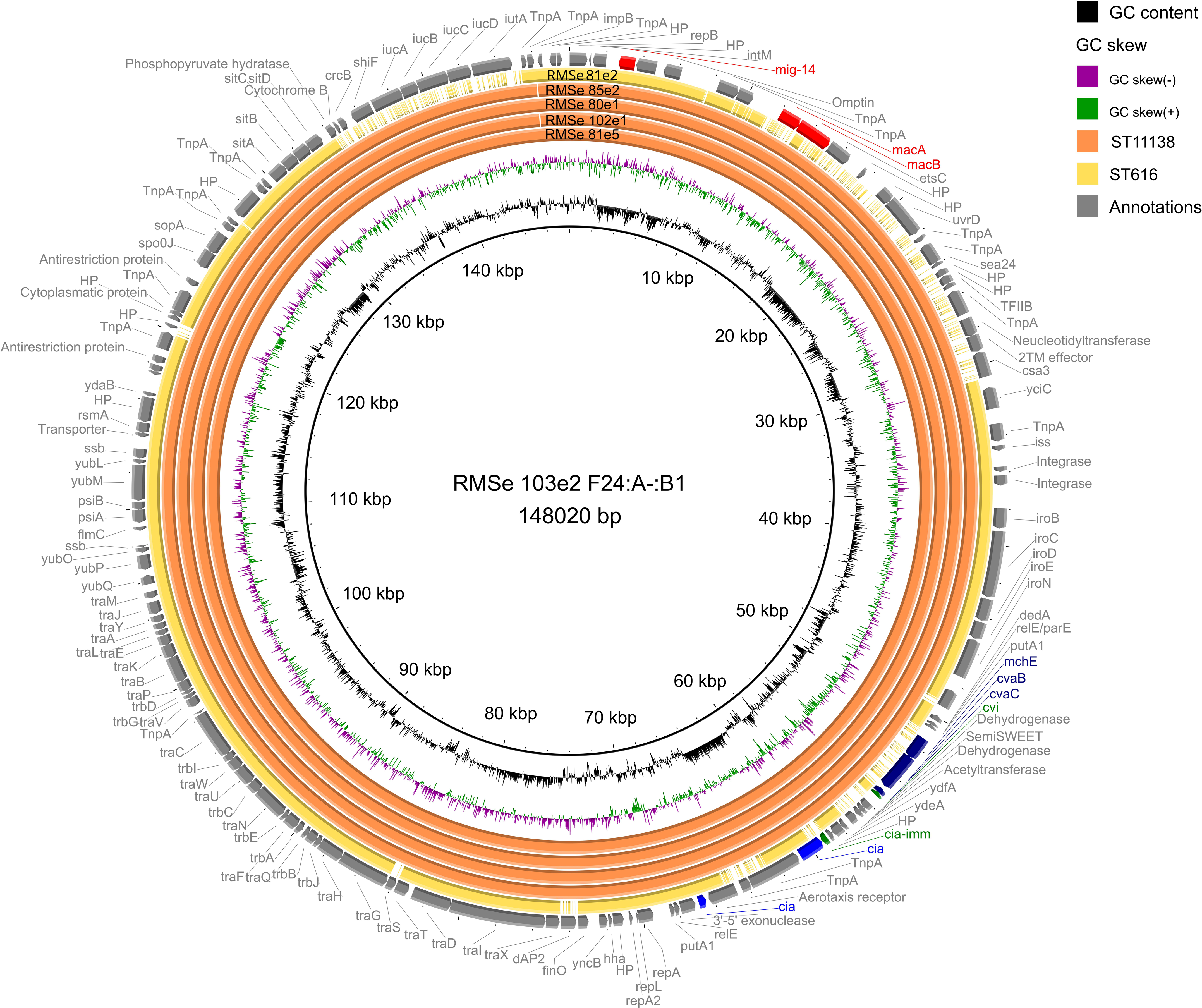
Alignment of the F24:A-:B1 plasmids from selected ST11138 and ST616 isolates. Colicin genes (blue), colicin immunity genes (green), and antibiotic resistance genes (red) are highlighted. Lighter shading indicates disrupted genes. Colour assignment of STs is shown in the legend. The F24:A-:B1 plasmid obtained from long-read sequencing of isolate RMSe 103e2 was used as the reference.

The F34:A-:B-plasmid, found exclusively in ST11893, exhibited high sequence uniformity across all isolates. In each case, the *cba* colicin gene was disrupted by the *bla*_CMY-2_ resistance gene. A previously collected ST11893 isolate from an earlier study^5^ was included for comparison and displayed 100% sequence identity (Supplementary figure S4). Among all analysed plasmids, only one isolate differed by a single SNP; the remaining sequences were identical (Supplementary table S4).

The IncI1/ST3 plasmid alignment included representatives from all host STs in this study, as well as ST23 and ST540 sequences from the previous dataset^5^. Plasmids from ST117, ST165, ST919, and ST11138 exhibited 100% sequence identity (Supplementary figure S5). In contrast, plasmids from ST616, ST665, ST23, and ST540 showed variation, particularly in transposase-associated regions. SNP analysis of the IncI1/ST3 revealed high overall uniformity, with an average of 2.85 SNPs across all STs (Supplementary table S4). The IncI1/ST3 plasmid also carried complete *bla*_CTX-M-1_ gene for ESBL production in all STs. Due to its carriage of the *cib* colicin gene, absent from all other plasmids, the IncI1/ST3 plasmid was originally thought to be the main driver of the varying strain persistence.

## Discussion

In this study, we provide evidence that Caspian gulls are capable of maintaining cefotaxime-resistant *E. coli* strains in their gut for at least three months, continuously shedding ESBL-producing bacteria into the environment and serving as a secondary source for other birds in the flock. This is the longest longitudinal study to date investigating carriage of antibiotic-resistant bacteria in the gut microbiome of wild migratory birds, offering unprecedented insights into the persistence and dynamics of resistance in a natural host over time. The study resulted also in the longest documented duration of resistant bacteria carriage in wild birds, as previous studies have reported gulls shedding colistin-resistant *E. coli* for up to 16 days, after which the bacteria were no longer detected despite continued monitoring^16^, and mallards harbouring ESBL-producing *E. coli* for 29 days, with the experiment ending at that point^14^. At the time of the initial sampling at the gulls’ breeding site, only three out of five individuals carried resistant *E. coli*, with ST11893 predominantly present in all three and ST665, harbouring the IncI1/ST3 plasmid, found in a single gull. ST11893 was also previously identified as the most prevalent resistant *E. coli* strain in the same gull colony, where it was detected across multiple individuals in a single time point sampling^5^. By week 4, six different STs carrying the IncI1/ST3, likely due to its previously reported ability to disseminate efficiently^49^, began to appear, with ST11138 rapidly outcompeting all other strains. ST11893 disappeared by the second sampling, and no other STs lacking the IncI1/ST3 plasmid were detected afterwards. Although overall colonization rates declined by the end of the experiment, shedding was still observed along with ongoing transmission of strains between individual gulls, indicating a continued risk of environmental dissemination and spread to other hosts. Unlike *in vitro* experiments or *in silico* models, our study tracked the persistence and transmission of resistant bacteria in live wild birds under biologically relevant conditions, providing ecologically grounded evidence of strain dynamics and plasmid spread.

Due to the observed dominance of a single ST, four STs (ST11893, ST11138, ST165, ST665) were selected for further analyses to investigate the mechanisms behind this phenomenon. Competition experiments between ST11893 and ST11138 revealed a clear dominance of the latter, with nearly 97% of colonies belonging to ST11138. Although plasmid transfer of IncF and IncI1 was also detected, conjugation rates were low and unlikely to have influenced the outcome. The high fitness of ST11138 was evident not only in direct competition assays but also in fitness measurements, where it displayed significantly higher growth than all other tested STs. In contrast, ST665 and ST11893 exhibited the lowest fitness and, although present at the initial sampling, both were not detected in subsequent weeks. The IncI1/ST3 plasmid is generally considered to impose low, if any, fitness costs on its bacterial hosts^49,50^. A similar trend was observed in our study, where strain fitness did not appear to correlate with potential plasmid-associated fitness costs, as both the most and least successful STs carried the IncI1/ST3 plasmid.

None of the tested STs harboured IncF plasmids with identical RST, further highlighting that plasmid type alone cannot fully explain the observed differences in persistence. However, a more than 200-fold difference in conjugation frequency was observed, with the IncF plasmid transferring at much lower rates than the IncI1/ST3 plasmid. Among STs, ST165 exhibited the highest efficiency in disseminating the IncI1/ST3 plasmid, followed by ST11138. Notably, these two STs were the only ones consistently detected in the gulls from week six onward. Genomic analyses suggested that colicin production could contribute to ST persistence, prompting us to perform assays focused on evaluating bacteriocin activity. As expected, ST11893 displayed no inhibitory activity against three indicator strains, suggesting a lack of production for three of the six colicins tested, as opposed to the other STs, which showed broader activity. This absence of bacteriocin production is likely a key factor contributing to the disappearance of ST11893. In contrast, ST11138 exhibited the highest colicin production, supporting its prevalence over the other STs.

The presence of colicin and colicin immunity genes was further investigated using long-read analysis, which allowed precise localization of these genes on individual plasmids. ST11893 was the only ST lacking the *cib* gene, which was located on the IncI1/ST3 plasmid in all other STs. This absence corresponds with observed reduced colicin activity and likely contributed to the early disappearance of ST11893, which was outcompeted by various emergent STs with the IncI1/ST3 plasmid. Notably, the *cib* region was conserved across all isolates harbouring this plasmid regardless of ST, suggesting strong plasmid stability and horizontal dissemination. Although ST11893 was the only ST carrying a complete *cba* gene, which could have potentially served as a competitive mechanism, it was located on an F34:A-:B-plasmid and disrupted by the *bla*_CMY-2_ gene and therefore did not provide the strain with any evolutionary advantage. The same plasmid, including the disruption, was found in ST11893 isolates collected from multiple birds during a previous study at the same gull colony^5^. The gulls analysed in our study were, however, physically separated from the colony throughout the study, indicating that the strain with the exact plasmid managed to persist in the bacteria even with no outside contact with the colony, which could otherwise serve as a secondary source. While the F34:A-:B-plasmid also uniquely harboured a *cma* gene region, its associated colicin activity was insufficient to support persistence of ST11893 in the presence of more competitive STs. The key determinant of persistence appeared to be the F24:A-:B1 plasmid, carried by all ST11138 isolates and a single ST616 isolate. This plasmid encoded the *cvaC* gene, which was absent or incomplete in all other plasmids and STs. The colicin gene region on F24:A-:B1 was identical across all ST11138 isolates, while the plasmid sequence in the ST616 isolate showed notable divergence, likely explaining the dominance of ST11138. The second ST persisting alongside ST11138 until the end of the experiment was ST165, which carried an F-:A-:B73 plasmid. This plasmid harboured the *cma* gene and its corresponding immunity gene, similarly to the F34:A-:B-plasmid in ST11893. However, it appeared to confer an evolutionary advantage in ST165 due to its co-occurrence with the IncI1/ST3 plasmid carrying the *cib* gene region, which was absent in ST11893.

Together, these findings demonstrate that the persistence and competitive success of resistant *E. coli* strains in wild gulls are shaped by a complex interplay of plasmid-associated traits and ecological dynamics. While plasmid carriage alone did not correlate with strain fitness, the presence of specific colicin and colicin immunity genes emerged as a key factor influencing long-term colonization. Our study highlights the importance of bacteriocin-mediated interactions as a selective force within microbial communities and supports the view that mobile genetic elements contribute not only to AMR dissemination but also to competition among strains within wildlife reservoirs. Moreover, colicin production associated with AMR is particularly found in epidemic plasmid lineages found in multi-drug resistant bacteria. These conclusions are consistent with recent evidence showing that colonization history and host ecological context strongly influence the evolution of AMR in the gut microbiome^51^. Similarly to those findings, our results suggest that early colonizing strains equipped with strong competitive traits, such as colicin production, are more likely to persist and dominate, highlighting the importance of microbial community dynamics in shaping resistance patterns even in the absence of selective antibiotic pressure.

In summary, our study shows that the long-term persistence of cephalosporin-resistant *E. coli* in Caspian gulls is shaped not simply by plasmid carriage or initial prevalence, but by a multifaceted combination of bacterial fitness, plasmid dissemination efficiency, and competitive traits such as colicin production. These features enabled certain strains, notably ST11138 and ST165, to dominate the gut microbiome and sustain colonization over time, even without ongoing antibiotic selective pressure. By capturing these processes in live wild birds under naturalistic conditions and across an extended period, our study provides rare and ecologically grounded insights into how AMR is maintained outside clinical or laboratory settings. This unique host-based perspective shows that resistance dynamics in wildlife reservoirs are governed as much by microbial interactions and evolutionary persistence strategies as by the presence of ARGs themselves. Importantly, our findings underscore the ecological novelty of longitudinal *in vivo* tracking of AMR persistence in natural hosts and highlight the need to consider ecological and microbiome-level interactions when assessing the environmental dimension of AMR under the One Health concept.

## Data availability

All data from whole-genome sequencing are available at BioProject number PRJNA1144792.

## Acknowledgements

We thank Jarmila Lausova, Dana Cervinkova, Martina Masarikova, Tomas Nohejl, and Hassan Tarabai from University of Veterinary Sciences Brno (Czech Republic) for their help in the field and in the laboratory. We would also like to acknowledge the RECETOX Research Infrastructure for letting us utilize their Synergy HT reader for the fitness measurements.

## Authors’ contribution

MRu performed laboratory experiments, analysed and interpreted the data, created visualisations and wrote the manuscript. JP analysed the data, created visualisations and revised the manuscript. KN analysed and interpreted the data and revised the manuscript. Mry performed laboratory experiments and revised the manuscript. RP performed laboratory experiments, analysed the data and revised the manuscript. SK collected the data and revised the manuscript. IL supervised the work, acquired funding and revised the manuscript. MD supervised the work, designed the project, acquired funding and revised the manuscript.

## Ethics declarations

### Ethical approval

The collection of samples was part of a larger telemetry study, which was permitted by the local Czech Nature Protection Authorities (Permissions S-JMK78643/2018 OŽP/Ško and S-JMK 40970/2019 OŽP/Ško).

